# Beyond the Panglossian paradigm: adaptation, constraint, and anuran functional trait evolution

**DOI:** 10.64898/2026.07.20.739646

**Authors:** Menna Jones, Graham J. Slater

## Abstract

Recognizing patterns in functional trait evolution is a necessary step in testing macroevolutionary questions. Quantification of these patterns and interpretation of their generative processes relies on an ever-expanding suite of comparative approaches, but current methods oversimplify the process-to-pattern mapping. This simplification may promote binary classifications of patterns and their drivers, such as ‘adaptive versus non-adaptive’ or ‘constrained versus unconstrained’. A potentially more robust and evolutionarily informative alternative is to fit an expanded suite of evolutionary models at different levels of taxonomic or ecological resolution to reveal how evolutionary drivers imprint simultaneously on observed patterns of trait evolution. Here, we perform a clade-wide analysis of functional trait evolution across Anura (frogs and toads). Focusing on three functionally important traits, we quantify the relative fit of random, directional, and bounded models at the intra- and inter-microhabitat levels to explore how changes in trait function across microhabitats translate into different evolutionary regimes. We recover heterogeneous regime-level distributions of model support that imply complex underlying evolutionary dynamics, while also revealing methodological biases and model identifiability issues. These findings underscore the need to develop more robust tools for evolutionary model fitting and advance beyond binary frameworks for interpreting evolutionary processes.

## Introduction

Comparative studies are crucial tools for recognizing patterns in trait data and for inferring the nature of the evolutionary processes that generated them. The earliest comparative studies of macroevolutionary pattern and process were strictly empirical studies of the fossil record. For example, the Cenozoic record of North American mammals shows independent increases in average and maximum body size, suggesting a more general tendency for lineages to increase in size through their phylogenetic history (“Cope’s rule” Cope, 1887). Similarly, Eldredge and Gould used the absence of morphologically intermediate fossils as evidence for rapid phenotypic evolution in small, peripherally isolated populations (Grant, 1963; Mayr, 1942; Simpson, 1944), yielding a pattern that they termed “punctuated equilibrium” (Eldredge and Gould, 1972) and which contrasted the gradualistic view of attributing gaps to fossil record incompleteness (Mayr, 1963). The subsequent utilization of stochastic simulation studies in paleobiology (Raup et al., 1973) represented a shift towards what we recognize as modern-day comparative methods. The introduction of phylogenetic independent contrasts (PICs: Felsenstein, 1985) formally established the field of Phylogenetic Comparative Methods (PCMs), which now stretches far beyond fitting statistical Brownian motion models to patterns in trait data (Felsenstein, 1973, 1985), to acknowledge patterns that appear directional (Butler and King, 2004; Hansen, 1997; Harmon et al., 2010; Hunt, 2007; Hunt et al., 2015; Lande, 1976; Pagel, 1999), and vary across space, time, and taxonomic and/or ecological groups (Beaulieu et al., 2012; Eastman et al., 2011; Ingram and Mahler, 2013; Martin et al., 2023; O’Meara et al., 2006; Rabosky, 2009; Revell and Collar, 2009; Revell et al., 2024; Slater et al., 2010; Thomas et al., 2006; Uyeda and Harmon, 2014). Although these approaches have substantially advanced our understanding of the tempo and mode in phenotypic evolution over geological timescales, they are imperfect tools. Paleobiologists have long recognized that distinct evolutionary processes can leave the same patterns in comparative data derived from the fossil record (Foote, 1996; Raup and Gould, 1974), and this challenge is increasingly apparent to practitioners of phylogenetic comparative methods Uyeda et al. (2015). The questions of where these conflations reside and how to mitigate their effects remains poorly understood, however.

The most complex evolutionary patterns occur at megaevolutionary levels (Cooney et al., 2017; Simpson, 1944), but these are likely emergent properties of diverse processes that occur at lower levels of the phylogenetic hierarchy (Jablonski, 2007). For example, natural selection acts at the organismal level to exploit variation between individuals and drive mean trait values in a population towards optimal values. These optima confer maximum fitness to individuals via their effect on organismal performance (Arnold, 1983) and generate a predictable shift in trait values from ancestor to descendant while constraining the observed range of phenotypes. Lande (1976) developed a quantitative genetic model to describe this process which Hansen (1997) later noted corresponds exactly to an Ornstein-Uhlenbeck (OU) diffusion process and, because directional patterns in species-mean trait values across phylogeny are also viewed as evidence for underlying adaptive processes, he developed a macroevolutionary OU model that accounts for species’ covariances due to shared ancestry and which, via its many extensions (Beaulieu et al., 2012; Butler and King, 2004; Uyeda and Harmon, 2014) can be used to map the evolution of the adaptive landscape itself (Arnold et al., 2001; Simpson, 1944, 1953). However, selection does not act in isolation; constraints on trait evolution are present at all levels of the evolutionary hierarchy and range in form from developmental (Alberch, 1989; Gould and Lewontin, 1979; Hanken and Wake, 1993; Hopkins and Smith, 2015; Liu and Robinson-Rechavi, 2018; Maynard Smith et al., 1985; Wake, 1982), to physiological (Kleiber, 1947;

Qu and Wiens, 2020; Rayner, 1988; Schmidt-Nielsen, 1984; von May et al., 2017), to ecological (Berven et al., 1979; Graham et al., 1995; Losos, 2010; Martof, 1956; Rabosky, 2009; Slater, 2013; Vermeij, 1987). Yet, in the field of phylogenetic comparative biology, the evolution of organismal form, function, and behavior is still largely attributed to adaptation and selection alone, reinforcing a historical Panglossian paradigm (Gould and Lewontin, 1979) that fails to acknowledge the complex interplay between the adaptive and constraining forces that mold the phenotype (Losos, 2011). This bias is reflected in the assumptions underlying many phylogenetic comparative methods. For example, (Boyko and Rabosky, 2026) recently criticized the common practice of representing macroevolution as occurring within a Euclidean trait space because this geometry oversimplifies phenotypic evolution and poorly represents the influence of complex developmental and genetic constraints. This can result, though unintentionally, in a bias towards favoring simpler, more homogeneous explanations for evolutionary patterns, many of which center upon adaptation and selection (Beaulieu et al., 2012; Butler and King, 2004; Uyeda and Harmon, 2014). These methodological choices directly shape how evolutionary processes are defined and compared within model-based frameworks.

In this setting, no single model of trait evolution can adequately capture the interplay between adaptive and constraining forces. A typical comparative framework may involve comparing the fit of models of strictly adaptive evolution (an Ornstein-Uhlenbeck (OU) process Hansen, 1997; Lande, 1976) to that of a null model, most often an unbiased Brownian motion (BM: Felsenstein, 1973; Harvey and Pagel, 1991) in which trait variation accrues gradually over time along the branches of a phylogeny. However, this approach leads to an unnecessarily restrictive dichotomy in which low phylogenetic signal in comparative data is taken as evidence for dynamic adaptation, while high signal is frequently (though incorrectly) interpreted as evidence for drift. The implementation of a bounded BM model (Boucher and Démery, 2016) succeeds in allowing us to model several forms of constrained evolution that are not necessarily adaptive. Under one interpretation, bounded dynamics represent stochastic drift within fixed upper and lower limits, making the model particularly appropriate for traits such as body size, the evolution of which is constrained by environmental, developmental, and physiological factors across clades and taxonomic scales (Brown et al., 2004; Graham et al., 1995; Hanken and Wake, 1993; Harrison et al., 2010; Levy and Heald, 2016;

Peters, 1983; Rayner, 1988; Schmidt-Nielsen, 1984; Wake, 1982). However, analogous to how Brownian motion can arise from fluctuating selection around a moving adaptive optimum (Hansen and Martins, 1996; Lande, 1976), bounded BM may also represent evolution on a dynamic adaptive landscape in which adaptive peaks shift through time while remaining constrained by organismal limits (Boucher and Démery, 2016). In this scenario, bounded BM captures the joint influence of adaptation and intrinsic constraints on phenotypic evolution. In either interpretation, provided they are realized, the imposed bounds ultimately govern evolutionary outcomes: selection can influence trajectories only within the bounded trait space, making bounded BM fundamentally a model of constraint.

Despite its relevance to such evolutionary scenarios, the bounded BM model is infrequently used, despite the fact that its omission may inflate apparent support for adaptive models due to the low phylogenetic signal and constrained trait variance characteristic of both processes (Foote, 1996; Revell et al., 2008). Indeed, several authors have noted that estimated model parameters from best-fitting OU models should be carefully evaluated to ensure that they are consistent with evidence for directional selection, rather than just noisy dynamics or data (Boettiger et al., 2012; Cressler et al., 2015; Pennell et al., 2015; Rabosky and Goldberg, 2015). One possible reason for the slow adoption of bounded models in studies that evaluate adaptive hypotheses for trait evolution is that there is currently no bounded equivalent to multi-regime OU model, which restricts the fitting of bounded models to single-regime settings. A potential solution that has yet to be explored in the literature, as far as we are aware, is to conduct analyses at both single- and multi-regime resolutions and then evaluate the congruence of both sets of results (c.f. Foote, 1996; O’Meara et al., 2006). This approach would enable relative evaluation of the (non-)adaptive constraining forces that shape trait evolution within regimes to across a clade, provided that the two models are distinguishable in the focal clades.

In this study, we take exactly this approach to evaluate competing hypotheses regarding the macroevolutionary driving forces of functional trait evolution, focusing on the order Anura (frogs and toads). Anura is a popular target for studies of ecomorphology and functional trait evolution (Emerson, 1976, 1979; Engelkes et al., 2020; Moen, 2019; Moen et al., 2013, 2016; Womack and Bell, 2020) in part due to its exceptional breadth along all axes of diversity (taxonomic, morphological, developmental, ecological, geographical). The estimated 7956 extant species are distributed across every continent bar Antarctica (AmphibiaWeb, 2026), with a notable concentration of taxonomic diversity in the Neotropics (44%, (Duellman, 1988)). The most well-cited evidence for adaptively-driven functional and morphological evolution in any clade is the recognition of similar (*ecomorphs*: Losos, 1990; Losos et al., 1998) across continents. In anurans, these ecomorphs have been repeatedly converged upon throughout clade history (Duellman, 1994; Moen et al., 2013, 2016), reflecting in part microhabitat-specific selection for certain trait functions (Harvey and Pagel, 1991; Losos, 2011; Moen et al., 2013, 2016; Simpson, 1953; Stepanova and Womack, 2020). For example, arboreality is associated with dilated digit pads and ‘toe disc apparatus’ (Emerson and Diehl, 1980) for surface adhesion (Emerson and Diehl, 1980; Ernst, 1973; Green, 1979; Hildebrand et al., 2001; Noble and Jaeckle, 1928), with slender bodies for weight distribution (Tyler, 1998; Zimkus et al., 2012). Burrowers, to contrast, have round bodies and squat limbs (Emerson, 1976, 1985, 1991; Gomes et al., 2009; Moen, 2019; Zug and Altig, 1978), with a shortened tibiofibula promoting forceful scooping of the substrate (Emerson, 1976) to facilitate efficient locomotion. Not all functional and performance-based traits exhibit clear differences across ecological regimes, however: body size range (Womack and Bell, 2020) and jump performances (Moen, 2019) are two examples of this. These findings cast doubt on a fully adaptationist view of anuran trait evolution and suggest the presence of overarching, clade-wide constraints. This is supported by both the clade-wide maintenance of the characteristic, ancestral bauplan (Lires et al., 2016; Wake, 1997) and the apparent bounds on observed ranges of certain functional traits such as body size (Womack and Bell, 2020).

To tease out the relative influences of constraint and adaptation on functional trait evolution across Anura, we take advantage of recent developments in comparative methods. We first compare the fit of a series of macroevolutionary models, including bounded Brownian motion and multi-regime Ornstein-Uhlenbeck models, across anuran phylogeny to determine the macroevolutionary hypotheses that emerge from a clade-level view. Motivated by the hypothesis that constraining and adaptive forces should leave different signatures in within-regime data, we then perform a series of analyses at the level of individual microhabitats to ask whether support for multi-regime adaptive models at the clade-level may actually be attributable to more heterogeneous dynamics. By utilizing recently developed models and both within and among regime analyses, we aim to facilitate interpretations of the evolutionary underpinnings of anuran functional trait evolution and point to new avenues for future model development.

## Materials & Methods

### Trait & microhabitat data

We focus on three functional traits that have previously been identified as important in anuran ecomorphology: relative hindlimb length, degree of toe webbing, and body size. To compute these metrics, we made use of the comprehensive, publicly available anuran trait database of Morinaga et al. (2023). These authors collected measurements from museum specimens representing 1234 species across 51 families which they combined with data from existing datasets (Mendoza et al., 2020; Moen et al., 2013, 2016, 2021).

Relative hindlimb length is expressed as a ratio between hindlimb length and body size (snout-vent length; SVL): 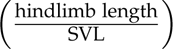. Anuran hindlimb morphology, in general, exhibits complex relationships with locomotor performance. Some comparative studies have found evidence for microhabitat-specific adaptive peaks in relative hindlimb length (Citadini et al., 2018; Moen, 2019; Stepanova and Womack, 2020), while others have indicated that this relationship holds only for burrowing species (Jorgensen and Reilly, 2013). Relatively longer hindlimbs improve swimming, jumping, and climbing performance (Citadini et al., 2018; Emerson, 1978; Moen et al., 2013; Nauwelaerts et al., 2007, 2005; Pérez-Ben et al., 2024; Petrović et al., 2017; Rebelo and Measey, 2019; Zug, 1972), while burrowing species boast short, squat hindlimbs that are optimized for forceful locomotion through the substrate (Citadini et al., 2018; Engelkes et al., 2020; Leavey et al., 2023). Based on previous work, we expect to recover a clear signal of dynamic adaptation in relative hindlimb length, present as strong support for microhabitat-specific optima and variance parameters. Toe webbing is quantified in different ways throughout the literature.

Past authors have based their metrics on both the total webbing area between hind limb digits (Moen and Wiens, 2017; Morinaga et al., 2023) and solely the webbed area between the fourth and fifth toes (e.g., Vidal-Garćıa and Keogh, 2015), typically standardized by body size (snout-vent length; SVL) to isolate differences in relative webbing between taxa. Discrete measurements are possible but less common, an example being the counts of webbing-free phalanges on toe IV (Zimkus et al., 2012). Here, we quantify toe webbing as the ratio of inter-digit webbing area to foot length 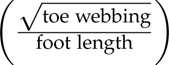. We acknowledge that our decision to standardize by foot length is unique; we do so because standardizing by body length makes the unnecessary and perhaps unreasonable assumption that toe webbing scales linearly with body size, independent of variation in toe or foot lengths. Thus, we consider that the functional importance of toe webbing can only be meaningful when considering the context of foot size. Toe webbing has functional relevance to both gliding in the arboreal environment and swimming in aquatic environments (Goldberg and Fabrezi, 2008; Tyler, 1998; Zimkus et al., 2012), as well as stability in terrestrial environments (Zimkus et al., 2012), suggesting that signals of adaptation might reasonably be expected.

Body size, measured as snout-vent length (SVL) of an adult individual, ranges from approximately 6.5 mm (*Brachycephalus pulex* (Bolaños et al., 2024)) to 320 mm (*Conraua goliath*) across Anura (Womack and Bell, 2020). It is commonly employed as an ecological proxy linked to diverse ecological, morphological, performance-based, and physiological traits, such as jump distance (Emerson, 1978; Gomes et al., 2009; Zug, 1972; Zug and Altig, 1978), desiccation rate (Newman and Dunham, 1994), reproductive mode (Gomez-Mestre et al., 2012; Levy and Heald, 2016; Womack and Bell, 2020), and metabolic rate (Brown et al., 2004; White et al., 2006). Microhabitat-associated differences in body size have not been consistently recovered in comparative analyses (Citadini et al., 2018; Vidal-Garćıa et al., 2014; Womack and Bell, 2020), and the recognition of hard trait limits across Anura specifically (Womack and Bell, 2020) suggests a highly constrained evolutionary trajectory for body size in Anura, with minor influence from microhabitat-specific selection. This makes it an interesting functional trait to include in our study. We log-transformed the three morphological traits prior to all analysis. Values reported in the main text are back-transformed for ease of interpretation.

To evaluate whether functional trait evolution is driven by regime-specific adaptations, we focused on adult microhabitat, acknowledging that oviposition and mating may occur elsewhere. We used microhabitat classifications of Moen and Wiens (2017), which are defined using substrate-use patterns (Moen et al., 2016). Four main microhabitat categories are recognized: 1) aquatic (rarely leaves a body of water), 4) burrowing, 3) terrestrial (found on the ground, away from water), and 4) arboreal (resides in above-ground vegetation). Intermediate patterns of substrate use are also sometimes recognized, where a taxon uses two substrates in a sub-equal fashion. For this study we consider semi-arboreal as a terrestrial-arboreal combination and semi-aquatic as a terrestrial-aquatic combination. We removed microhabitats that were defined but barely represented in the dataset (aquatic-burrowing: *n* = 1; arboreal-burrowing: *n* = 2; burrowing-semi-arboreal: *n* = 6; semi-burrowing: *n* = 14; terrestrial-torrential:*n* = 3) to avoid issues with downstream statistical analyses. Finally, we treat torrential (living in small, rapidly flowing mountain streams) as a distinct microhabitat due to the unique challenges imposed by this near-arboreal-aquatic environment (Endlein et al., 2013). Although we recognize the shortfalls of discretizing an inherently continuous ecological spectrum, such classifications are standard practice in the literature, and developing a more refined scheme is beyond the scope of this study.

### Phylogeny

To investigate the evolution of the three functional traits in a phylogenetic context, the time-calibrated molecular phylogeny of 5424 amphibian species was taken from Portik et al. (2023) and pruned to the 927 species for which functional trait and microhabitat data were available (aquatic: *n* = 39; semi-aquatic *n* = 70; torrential: *n* = 48; burrowing: *n* = 44; terrestrial: *n* = 352; semi-arboreal: *n* = 77; arboreal: *n* = 297; Fig. 1).

**Figure 1:**
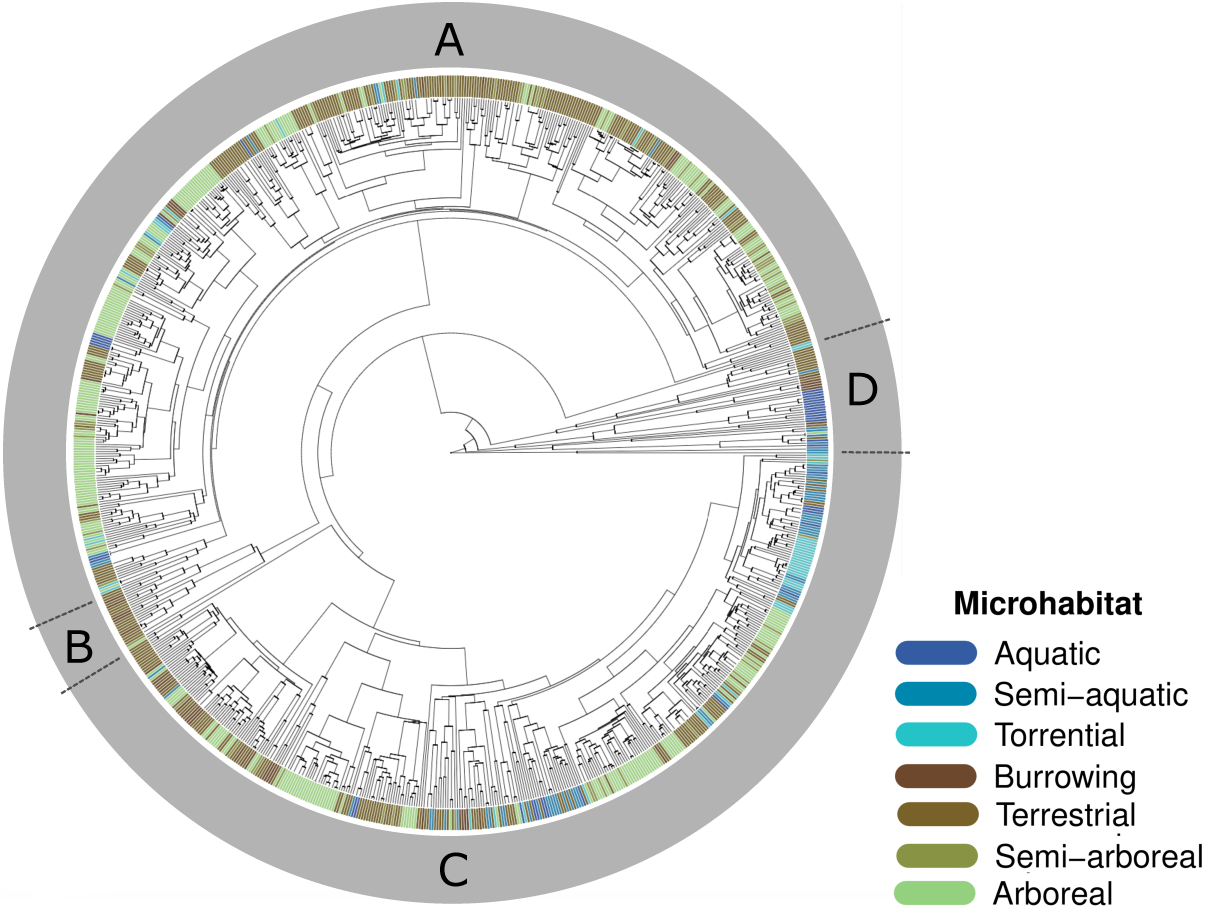
Time-calibrated phylogeny of 927 anuran species with tip colors representing assigned microhabitat. Major clades are indicated as A) Hyloidea, B) Myobatrachoidea, Sooglossidae, C) Ranoidea, D) Discoglos-soidea, Leiopelmatoidea, Pelobatoidea, Heleophrynidae and Pipoidea. The original phylogeny came from (Portik et al., 2023).

### Clade level view of anuran functional trait evolution

To evaluate support for different modes of functional trait evolution across anuran phylogeny, we fit eight models of continuous trait evolution to each set of functional trait data. Single-(BM1) and multi-regime (BMS) Brownian motion both reflect unconstrained diffusive processes defined by a root state (*x*_0_) and variance parameter(s) (*σ*^2^). Support for these processes is often interpreted as evidence for a random evolutionary trajectory that accumulates variance in proportion to time, though it should also be noted that they are consistent with adaptation chasing stochastically fluctuating optima (Revell et al., 2008). Bounded BM introduces constraint in the form of bounds that restrict the accrual of trait variance (Boucher and Démery, 2016). Support for bounded BM is consistent with non-directional, diffusive evolutionary dynamics that are constrained in a non-adaptive manner (Foote, 1996).

Single-regime OU (OU) is the simplest adaptive model considered, describing stabilizing selection of a given strength *α* towards an optimum (*θ*), the position of which is a product of selective and genetic constraint (Hansen, 1997). Multi-regime OU models are typically assessed when a trait is hypothesized to have evolved under different, ecologically defined regimes associated with distinct sets of selection (Butler and King, 2004). Thus, four multi-regime OU models were also considered in this study. The first estimates regime-specific optima for each microhabitat while keeping *α* and variance parameters (*σ*^2^) constant (OUM). Increasingly complex variants of the OUM model allow *σ*^2^ (OUMV), *α* (OUMA), or both (OUMVA) to vary among regimes (Beaulieu et al., 2012). Single-regime bounded Brownian motion was fit using the bounded bm() function in the phytools() library (Revell, 2024), and the seven remaining models were fit with the OUwie() function in the OUwielibrary (Beaulieu and O’Meara, 2022) in R v. 4.4.1 (R Core Team, 2024). Relative model fits were evaluated using AIC weights (Wagenmakers and Farrell, 2004), where values closer to one indicate higher support for a given model relative to other models in the candidate set. Interpreting these weights should be done cautiously and whilst acknowledging that the true model may not be in the candidate set, which risks recovery of a best-fitting model that is not a representation of the true generating process.

Multi-regime models require that the evolutionary history of the regimes also be known such that parameters can be optimized along internal branches of the phylogeny. Therefore, we performed a Maximum Likelihood (ML) ancestral state estimation for microhabitat data. We first fit equal rates (ER), symmetric rates (SYM), and all-rates-different (ARD) models to our phylogeny and discrete tip data; however, only the ER successfully converged upon a solution, regardless of functions used (ape :: ace() (Paradis et al., 2004), phytools :: make.simmap() (Revell, 2024)). The failure of SYM and ARD models to converge reflects the high number of parameters associated with these models. Whilst intermediate, less parameter-rich models are possible, ancestral state estimation is not the focus of this study. Thus, we proceeded with the ER estimates of ancestral microhabitat regimes assigned the most likely estimated state under this model to each internal node to produce a fully mapped phylogeny.

### Is support for multi-regime adaptive models at the clade level attributable to heterogeneous dynamics at the regime level?

High support for multi-regime OU models may reflect dynamic adaptation but, as OU and bounded models share low phylogenetic signal, such support may also be consistent with constrained evolutionary dynamics, the nature of which vary by evolutionary regime. There is currently no multi-regime bounded model against which to directly compare the fit of a multi-regime OU model. We therefore performed a pruning procedure to differentiate between these two evolutionary scenarios.

Our approach is motivated by the prediction that if OUM is both the best-fitting and true model for the complete anuran dataset, then extracting regime-specific sub-trees and refitting single regime models to each individually should yield complementary support for OU1 in each case. The logic behind this approach may not be clear to all readers but we assert that it is a reasonable solution to an otherwise unresolvable (at present) problem. Consider, for example, a reciprocally monophyletic pair of clades evolving under distinct adaptive regimes. In such a case, support for a flavor of multi-regime OU model (Beaulieu et al., 2012; Butler and King, 2004) over BM-based alternatives could be decomposed to a set of tests within each of the clades in which we would expect to recover support for single peak OU models over BM-based models. This same logic forms the basis of the “censored” tests of rate heterogeneity employed by O’Meara et al. (2006). If, instead, this procedure yields support for a bounded model, then support for OUM at the level of Anura may be interpreted as erroneously resulting from low phylogenetic signal and a lack of appropriate multi-regime bounded models against which to directly evaluate. Our pruning approach therefore provides a useful way to identify potential conflation between adaptive and constrained dynamics that cannot be recognized from clade-wide analyses alone. In each case, relative model support was evaluated using AIC weights.

A potential limitation of this approach is that pruning regime-specific sub-trees can distort internal branch lengths if the ecological regimes are not monophyletic groups (c.f. O’Meara et al., 2006). This means that extraction of regime-specific sub-trees increases the total branch length over which the evolutionary model is fitted, relative to the mapped clade-level tree, by including branches that potential evolved under alternate evolutionary modes. A second, more general concern is parameter identifiability in small phylogenies. In particular, support for simpler models such as Brownian motion may reflect limited statistical power rather than true absence of alternative processes when clade sizes fall below some threshold size (Boettiger et al., 2012). Furthermore, this identifiability issue is likely to be particularly strongly felt when *α* is small relative to tree depth, resulting in genuine but weak mean-reversion in the data (Boettiger et al., 2012). This limitation is not solely a finite-sample issue, but an intrinsic property of the OU model itself (Ho and Ané, 2014). One shared consequence of this pruning procedure could therefore be to artificially decrease support for a true OU model, in favor of unbounded or bounded Brownian motion. To assess whether this potential distortion invalidates our approach, we therefore performed a simulation study. We first extracted the ML parameter estimates associated with the most complex OUMV candidate models on a per-trait basis and used these to simulate data, prune out the sub-trees, and compare the fit of BM1, OU1, and bounded BM models to each, a procedure which was repeated 100 times. Assuming that the pruning procedure does not distort the data, OU1 should be recovered as the best supported model for each sub-tree dataset. Conversely, if we recover support for bounded BM or BM for each regime, this would indicate that the method manipulates internal topologies and biases model comparison away from the true model. Intermediate results may suggest issues with model identifiability relating to factors such as generating parameter values and/or clade size.

High error rates may be expected when comparing models that produce similar trait distributions, particularly when sample sizes are small (Boettiger et al., 2012; Cooper et al., 2016; Pennell et al., 2012), as is often the case for individual evolutionary regimes. Thus, even if our pruning method avoids distorting model-fitting results, difficulty in choosing between model classes may persist. To quantify the type I error rates associated with both OU and bounded generating models, we performed a further basic simulation study. We generated 100 independent trait datasets under an OU1 model, varying *σ*^2^ ∈ {0.01, 0.1, 1} and *α*, with *α* spanning a range of phylogenetic half-life values *T*_1/2_ ∈ {0.01, 0.05, 0.1, 0.5}. These values were chosen to minimize the identifiability problems associated with small *α* parameters (Boettiger et al., 2012; Ho and Ané, 2014), thus restricting our analysis to scenarios in which misclassification is unlikely to arise simply from OU models being statistically indistinguishable from Brownian motion. We then fit both OU and bounded models to each dataset and evaluated model support using AIC weights. We then repeated this procedure using a bounded generating model, varying *σ*^2^ to produce *τ* values equivalent to the specified *T*_1/2_ for the chosen bound breadth, {−1, 1}. This workflow yielded 18 generative models (12 × OU, 6 × bounded), each simulated across four tree sizes *n* ∈ {20, 50, 100, 200}.

## Results

### Ancestral microhabitat regime estimation

Ancestral state estimation under a model of equal transition rates between microhabitat pairs (ER) yielded a most likely ancestral state of terrestriality for Anura (Fig. S1) and a total of 299 transitions between microhabitat states averaged across 100 simulations (Table. S1). The most common transition was from terrestrial to arboreal (*n* = 72.3), followed by arboreal to terrestrial (*n* = 64.4) and terrestrial to semi-arboreal (*n* = 34.4). Of the estimated 555 transitions, 201 are out of the terrestrial state to others, contrasting the mere 30 transitions out of the aquatic state. All 42 possible transitions between states are inferred to have occurred.

### Clade-level view of anuran functional trait evolution

The OUMV model received the greatest support for describing how relative hindlimb length (*W_OUMV_* = 1), toe webbing (*W_OUMV_* = 1), and body size (*W_OUMV_* = 1) evolved across the anuran phylogeny (Fig. 2). For relative hindlimb length, evolutionary rates are highest in the burrowing realm (*σ*^2^ = 1.3 × 10^−3^) and lowest in the semi-aquatic realm (*σ*^2^ = 3.3 × 10^−4^). The shared *α* parameter across regimes is estimated as 2.3 × 10^−2^ which corresponds to a phylogenetic half life of 30.5Myr, notably shorter than estimates recovered for the other traits. ML estimates of hindlimb optima vary notably between microhabitats (Fig. 3A): the lowest optimum is recovered in the burrowing realm (*θ* = 1.0), and the highest in the torrential realm (*θ* = 1.6). Despite the noticeable variation, standard error intervals surrounding five of the optima are overlapping and it is only in the torrential and burrowing realms that values are distinct.

**Figure 2:**
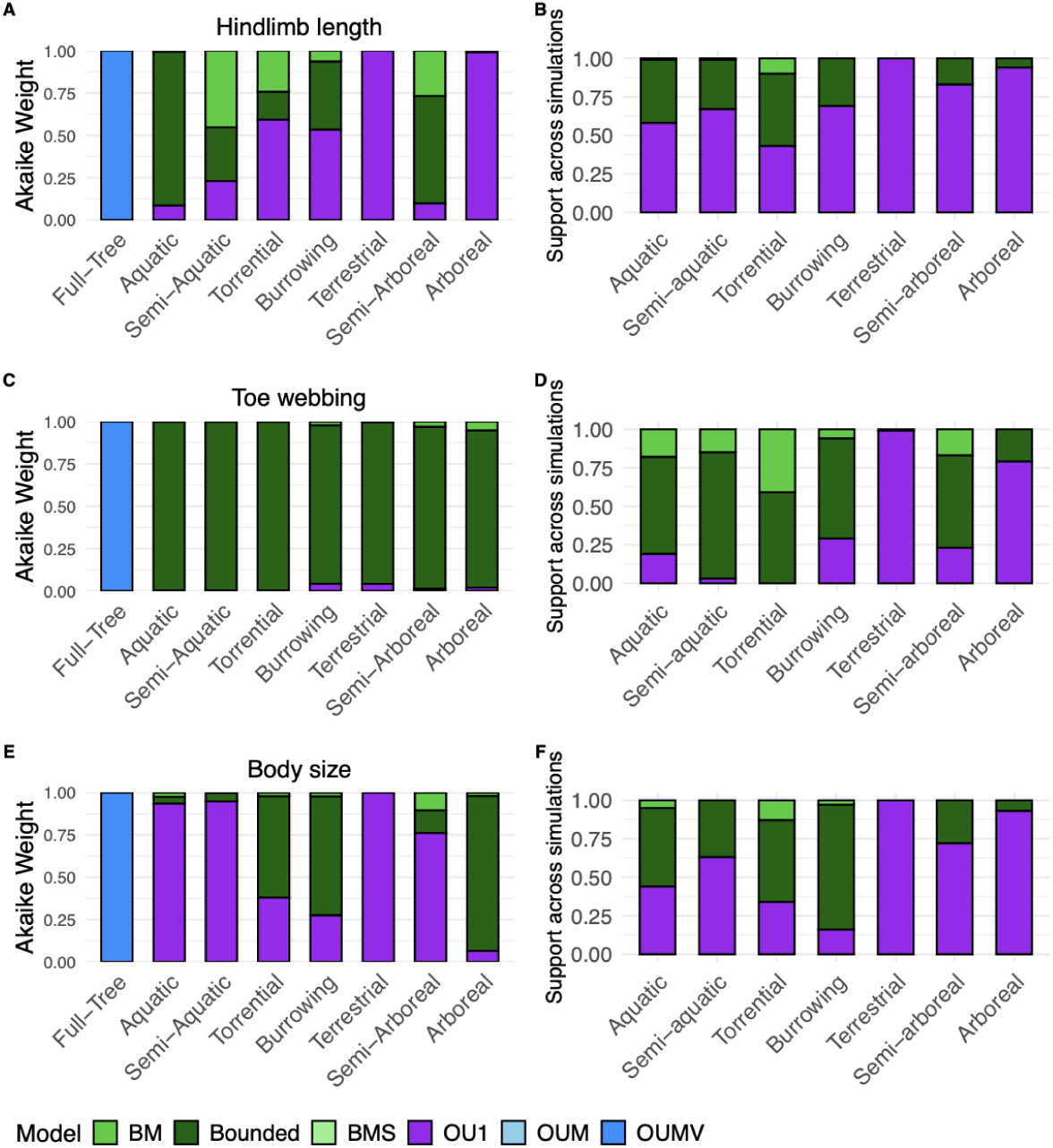
Left: evolution of relative hindlimb length (A), toe webbing (B) and body size (C) across the full tree and individual microhabitats. Each color represents a different candidate model. Support is calculated and plotted as AIC weights. Right: panels (B), (D), and (F) show the results of simulating across the tree under the ML estimates of OUMV parameters. Each bar shows the proportion by which each candidate model was favored across the 100 repeated simulations.

**Figure 3:**
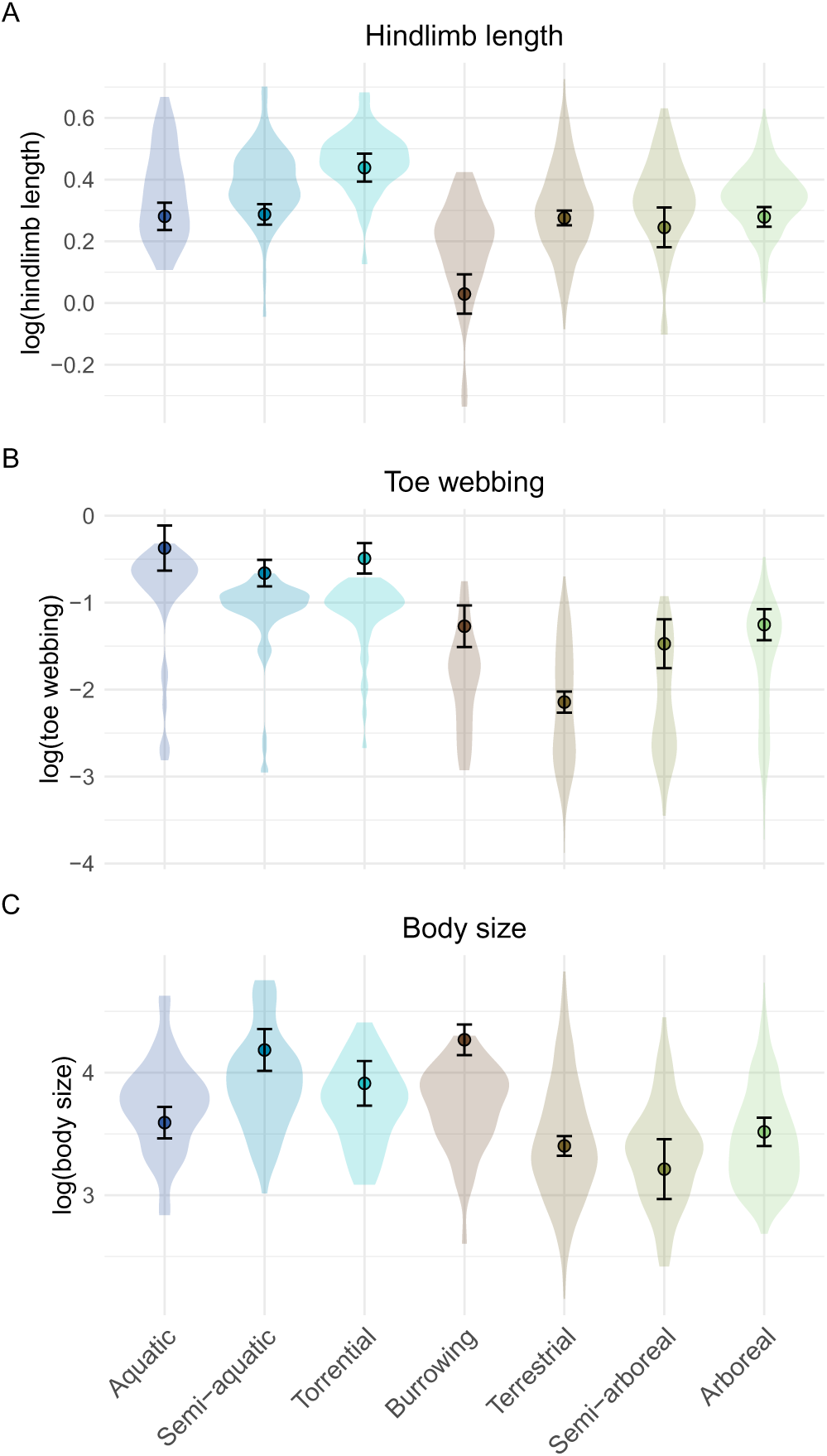
Log-transformed empirical distributions of relative hindlimb length (A), toe webbing (B), and body size (C) across individual microhabitats. Estimated relative hindlimb length (A), toe webbing (B), and body size (C) optima recovered from the ML OUMV model fit across the clade. Error bars are showing standard error (SE) and the background violin plots show the observed distribution in trait values post-log-transformation.

Optimal toe webbing values are greatest in the aquatic (*θ* = 0.69), torrential (*θ* = 0.61), and semi-aquatic (*θ* = 0.60) realms, and smallest in the terrestrial realm (*θ* = 0.12) (Fig. 3B). Rate estimates vary from 8.6 × 10^−4^ in the torrential realm to 1.4 × 10^−2^ in the terrestrial realm, with a shared alpha estimate of 1.7 × 10^−2^ (*T*_1/2_ = 41.4 Myr). Estimated optima are found to lie on the periphery of observed trait values in the (semi-)aquatic realms and even slightly above this range in the torrential realm. This is a stark contrast to the central position of estimated webbing optimum in the terrestrial realm which lies central in the relatively symmetrical distribution of observed values.

Patterns in body size evolutionary parameters show the largest optimum value in the burrowing realm (*θ* = 71.5 mm), which lies at the upper limit of the observed range of values (Fig. 3C). Evolutionary rate estimates are the lowest in this realm (*σ*^2^ = 2.0 × 10^−3^) and highest in the semi-aquatic realm (*σ*^2^ = 9.4 × 10^−3^). We estimate *α* = 2.0 × 10^−2^ (*T*_1/2_ = 34.6 Myr) for body size, which is intermediate between estimates for hindlimb length and toe webbing. Similar to the hindlimb length results, optima are overlapping between microhabitats which prevents diagnosing microhabitat-specific values.

### Is support for multi-regime adaptive models at the clade-level attributable to heterogeneous dynamics at the regime-level?

It is possible that support for multi-regime OU models with slow rates of adaptation reflect complex bounded dynamics, rather than adaptive evolution, for our three traits. To test this hypothesis in the absence of a multi-regime bounded model, we extracted comparative datasets representing each regime and compared the fit of three simple models, unbounded BM, bounded BM, and OU1, with the prediction that a true OUM(V) model across regimes would manifest in high support for the OU1 model within individual regimes. Regime-specific analyses of all three traits produced heterogeneous results (Fig. 2, Table S2) both within and between microhabitat regimes. For relative hindlimb length, some regimes (torrential, terrestrial, arboreal) yielded support for a best-fitting OU1 model, consistent with our initial predictions, but others (aquatic, burrowing, semi-arboreal) were best explained by a bounded Brownian motion model. BM received the least support overall, being favored only in the semi-aquatic realm (*W* = 0.497). Toe webbing results were more homogeneous: the bounded model received the highest support across all microhabitats, despite the support for clade-wide OUMV dynamics. Finally, body size evolution was best modeled with an OU1 model in the aquatic, semi-aquatic, terrestrial, and semi-arboreal regimes, and a bounded model in the torrential, burrowing, and arboreal regimes.

The pruning method used to isolate regimes was validated via simulations under the ML parameterization of the best-fitting clade-wide OUMV model for each trait. Bounded, BM, and OU1 models were subsequently fit to each regime to evaluate whether OU1 is preferentially supported given the overarching OUMV generating process being known. The results demonstrate, for each trait, instances of the bounded model receiving erroneously high support for within-regime dynamics (Table 1, Fig. 2). This is especially apparent for regimes containing fewer taxa (e.g. toe webbing in a semi-aquatic setting, body size in a burrowing setting). OU1 support was strongest in the most species-rich regimes, ranging across traits from *W_OU_*_1_ = 0.96 − 1 and *W_OU_*_1_ = 0.79 − 0.93 in the terrestrial and arboreal regimes, respectively. The ability to detect OU1 in the largest regimes suggests that support for a bounded model may result not from pruning method flaws but from limitations of comparative model fitting at small sample sizes when different models generate statistically similar tip data. The Spearman rank correlation between clade size and proportional support for OU1 model (Spearman, 1987) indicates a significant, positive linear relationship for body size (*ρ* = 0.964, *p <* 0.01), and positive but non-significant relationships for hindlimb length (*ρ* = 0.75, *p* = 0.052) and toe webbing (*ρ* = 0.536, *p* = 0.236).

**Table 1:** Results of the pruning method validation. For each trait, continuous data was simulated under the best-fitting OUMV model (using estimated parameters) across 100 simulations. Each microhabitat was subsequently pruned and BM, bounded, and OU1 models were fit. The number of tips in the sub-tree and the proportion of simulations in which each model received the greatest support (according to AIC weights) are recorded. The most frequently favored model in each set is given in bold.

| Microhabitat | *N* taxa | Hindlimb length |  |  | Toe webbing |  |  | Body size |  |  |
| --- | --- | --- | --- | --- | --- | --- | --- | --- | --- | --- |
|  |  | BM | Bounded | OU1 | BM | Bounded | OU1 | BM | Bounded | OU1 |
| Aquatic | 39 | 0.040 | 0.44 | **0.52** | 0.18 | **0.63** | 0.19 | 0.05 | **0.51** | 0.44 |
| Semi-aquatic | 70 | 0.084 | 0.37 | **0.54** | 0.15 | **0.82** | 0.03 | 0.00 | 0.37 | **0.63** |
| Torrential | 48 | 0.22 | **0.50** | 0.28 | 0.41 | **0.59** | 0.00 | 0.13 | **0.53** | 0.34 |
| Burrowing | 33 | 0.043 | 0.34 | **0.63** | 0.06 | **0.65** | 0.29 | 0.03 | **0.81** | 0.16 |
| Terrestrial | 352 | 0.00 | 0.00 | **1.00** | 0.00 | 0.01 | **0.99** | 0.00 | 0.00 | **1.00** |
| Semi-arboreal | 77 | 0.063 | 0.28 | **0.66** | 0.17 | **0.60** | 0.23 | 0.00 | 0.28 | **0.72** |
| Arboreal | 297 | 0.014 | 0.093 | **0.93** | 0.00 | 0.21 | **0.79** | 0.00 | 0.07 | **0.93** |

To disentangle whether identifiability issues observed in multi-regime and pruned analyses originate from the pruning procedure or from more fundamental limitations of model fitting, we performed an additional simulation study wherein we simulated under both OU1 and bounded BM over a range of tree sizes and model parameters and evaluated our ability to detect the true generating model using AIC weights. For a true OU1 model, its support hinges heavily on tree size, more so than parameter choice (Table 2). OU1 never outperforms the bounded alternative for the smallest tree size, and only does so for two model settings for a tree size of 50 (*T*_1/2_ = 0.1, *σ*^2^ = 0.1, 1). Counterintuitively, increasing *T*_1/2_ was found to increase OU1 support across tree sizes, yet this reverses when shifting from *T*_1/2_ = 0.1 to *T*_1/2_ = 0.5 and a decrease in OU1 support is recovered. Despite this, the OU1 model still garners majority support when *T*_1/2_ *>* 0.01 for tree sizes of 100 and above. In contrast, we recovered near-unanimous support for a bounded model across all tree sizes and simulation parameter combinations (Table 3): even at a tree size of 20, the minimum support for the true bounded model is 0.887. Thus, a real OU1 process may look like bounded evolution in small clades but bounded evolution tends to never look like adaptive evolution, regardless of clade size.

**Table 2:** OU1 simulation across a range of tree sizes and parameter combinations. Support values are averaged across 100 simulations. The favored model for each set of parameters is given in bold.

| $t_{1/2}$ | $\sigma^2$ | Tree size: 20 | | Tree size: 50 | | Tree size: 100 | | Tree size: 200 | |
| --- | --- | --- | --- | --- | --- | --- | --- | --- | --- |
|  |  | OU1 | Bounded | OU1 | Bounded | OU1 | Bounded | OU1 | Bounded |
| 0.01 | 0.01 | 0.107 | **0.893** | 0.125 | **0.875** | 0.223 | **0.777** | 0.395 | **0.605** |
| 0.01 | 0.1 | 0.105 | **0.895** | 0.157 | **0.843** | 0.239 | **0.761** | 0.398 | **0.602** |
| 0.01 | 1 | 0.113 | **0.887** | 0.127 | **0.873** | 0.180 | **0.820** | 0.306 | **0.694** |
| 0.05 | 0.01 | 0.141 | **0.859** | 0.396 | **0.604** | **0.699** | 0.301 | **0.978** | 0.022 |
| 0.05 | 0.1 | 0.152 | **0.848** | 0.398 | **0.602** | **0.663** | 0.337 | **0.945** | 0.055 |
| 0.05 | 1 | 0.182 | **0.818** | 0.337 | **0.663** | **0.723** | 0.277 | **0.923** | 0.077 |
| 0.1 | 0.01 | 0.223 | **0.777** | 0.496 | **0.504** | **0.876** | 0.124 | **0.998** | 0.002 |
| 0.1 | 0.1 | 0.162 | **0.838** | **0.525** | 0.475 | **0.882** | 0.118 | **0.999** | 0.001 |
| 0.1 | 1 | 0.244 | **0.756** | **0.526** | 0.474 | **0.855** | 0.145 | **0.998** | 0.002 |
| 0.5 | 0.01 | 0.210 | **0.790** | 0.403 | **0.597** | **0.629** | 0.371 | **0.804** | 0.196 |
| 0.5 | 0.1 | 0.212 | **0.788** | 0.406 | **0.594** | **0.549** | 0.451 | **0.782** | 0.218 |
| 0.5 | 1 | 0.209 | **0.791** | 0.386 | **0.614** | **0.596** | 0.404 | **0.820** | 0.180 |

**Table 3:** Bounded simulations across a range of tree sizes and parameter combinations. Support values are averaged across 100 simulations. The favored model for each set of parameters is given in bold.

| $\tau$ | $\sigma^2$ | Tree size: 20 | | Tree size: 50 | | Tree size: 100 | | Tree size: 200 | |
| --- | --- | --- | --- | --- | --- | --- | --- | --- | --- |
|  |  | OU1 | Bounded | OU1 | Bounded | OU1 | Bounded | OU1 | Bounded |
| 0.1 | 40 | 0.009 | **0.991** | 0.000 | **1.000** | 0.000 | **1.000** | 0.000 | **1.000** |
| 0.5 | 8 | 0.012 | **0.988** | 0.000 | **1.000** | 0.000 | **1.000** | 0.000 | **1.000** |
| 1 | 4 | 0.022 | **0.978** | 0.003 | **0.997** | 0.000 | **1.000** | 0.000 | **1.000** |
| 5 | 0.8 | 0.060 | **0.940** | 0.021 | **0.979** | 0.002 | **0.998** | 0.000 | **1.000** |
| 10 | 0.4 | 0.113 | **0.887** | 0.070 | **0.930** | 0.016 | **0.984** | 0.007 | **0.993** |
| 50 | 0.08 | 0.175 | **0.825** | 0.159 | **0.841** | 0.180 | **0.820** | 0.245 | **0.755** |

## Discussion

In recent years, the phylogenetic comparative toolkit has moved beyond the estimation of ancestral states and time-homogeneous rates of diffusive evolution for continuous traits (Felsenstein, 1973, 1985; O’Meara et al., 2006; Thomas et al., 2006) to also consider phenomenological models of adaptive radiation (Harmon et al., 2010) and process-based models of adaptive evolutionary dynamics (Beaulieu et al., 2012; Butler and King, 2004; Hansen, 1997). However, if comparative methods can be fooled by a many-to-one mapping of process to pattern (Foote, 1996; Revell et al., 2008), the insights we can gain into the processes underlying macroevolutionary patterns may be limited. This potential shortcoming is particularly concerning for cases where similar patterns in trait data are known to imply very distinct evolutionary modes, such as for adaptive and constraining forces (Losos, 2011). The issue is especially pertinent for the study of clades such as Anura, which exhibit extensive morphological, ecological, and reproductive diversity while remaining subject to constraints that differ in form and scope. For example, the architectural constraints imposed by the shared ancestral bauplan (Gould and Lewontin, 1979; Lires et al., 2016; Wake, 1997) act broadly across the clade, whereas certain ecological and morphological constraints vary with reproductive mode (Alberch, 1989; Duellman, 1994; Salthe and Duellman, 1973; Wake, 1982). We hypothesized that constraining and adaptive forces should leave distinct signatures in comparative data but that we are limited in our ability to approximate these processes by the models available to us, particularly when these dynamics vary across ecological evolutionary regimes. We tested this hypothesis by comparing the macroevolutionary interpretations that emerge from best-fitting models fitted to clade-wide and microhabitat-specific comparative datasets of three functional traits - relative hindlimb length, toe webbing, and body size - within Anura. Our results indicate that the clade-wide breadth in anuran diversity is generated by an accompanying breadth of evolutionary modes in functional trait evolution, determined in part by microhabitat regime. Evolutionary mode heterogeneity is suggestive of a confluence of constraining factors shaping functional trait evolution, yet this biological scenario is difficult to represent and interpret with the available comparative tools.

### Dangers of aligning with a Panglossian PCM Paradigm

Gould and Lewontin (1979) famously critiqued the tendency of biologists to see evidence of adaptation in every morphological feature and stressed the importance of considering constraint as a factor in organismal form. Yet, over 45 years later, the remnants of this Panglossian paradigm remain in the field of phylogenetic comparative methods as a simplified dichotomy between adaptive and non-adaptive modes of evolution, and statistical biases underlying the framework of comparative methods (Boyko and Rabosky, 2026). Our results suggest that, as Gould and Lewontin 1979 argued, this binary framework is not only insufficient to capture the multiple influences on functional trait evolution, but can also be positively misleading when applied to empirical systems.

Phylogenetic summary statistics are one tool employed to tease apart underlying evolutionary forces from statistical differences in the trait data they produce. Lande’s model of selection (Lande, 1976) describes how sufficiently strong selection will shift trait means toward fitness optima, typically interpreted in functional traits as peaks in organismal performance (Arnold, 1983). Lande’s model also implies that if the fitness peak is stable in time, then the long-term equilibrium process will be one of stable variance. Translated to the macroevolutionary case (Hansen, 1997), this process is one of low phylogenetic signal in which phylogenetic covariance cannot predict trait variables. Importantly, this same statistical pattern can arise under constraint-driven evolutionary dynamics. Low phylogenetic signal is therefore not diagnostic of adaptation alone, but is also consistent with bounded or constrained evolution (Revell et al., 2008). This ambiguity has been exploited in empirical work; for example, Hopkins and Smith (2015) attributed low signal and slow rates in post-Paleozoic regular echinoids to morphological constraint, contrasting them with more rapidly evolving irregular echinoids. But while this idea is mostly well-appreciated by comparative biologists, the fact that model-fitting is susceptible to the same problem if we fail to consider an appropriate suite of candidate models has not yet been adequately recognized. A lack, until recently, of appropriate models for describing bounded evolutionary processes led a few authors to leverage their similarity, in terms of pattern, to other, better understood models. For example, Slater (2013) used a single peak OU model to approximate constrained body size evolution in Mesozoic mammals relative to their Cenozoic descendants, with the rationale that both bounded Brownian motion and single stationary peak OU models should reduce the predictive value of the phylogenetic covariance matrix, relative to unbounded Brownian motion, and therefore result in a higher likelihood of observed data even after penalizing for the additional parameter. Although this approach allowed the rejection of a time-heterogeneous rate model in favor of a time-heterogeneous mode model (Slater, 2013) it ignored the statistical inconsistencies between the fitted model and the assumed bounded process and yielded estimated parameters that are uninterpretable in the context of the motivating hypotheses. Taken as a whole, these examples reinforce the failure of binary classifications in phylogenetic comparative biology and the need to explicitly test for a spectrum of influences on trait evolution.

The issue of model non-identifiability discussed above becomes apparent when interpreting the results from our clade-level analyses, which provide strong support for the role of selection in shaping anuran functional trait evolution. Under a strictly adaptationist interpretation, support for multi-regime OU models would be taken as evidence for distinct adaptive peaks, reflecting optimized functional performance of hindlimbs, toe webbing, and body size within particular ecological niches. However, patterns uncovered at the microhabitat resolution point to more heterogeneous dynamics that are not fully captured by this adaptive framework. Instead, signatures of constraint-driven evolution emerge, with multiple instances in which trait variation is limited not by attraction toward an optimum, but by bounds on the range of potential values. Even when selection is strong, these constraints constitute overarching limits on the extent to which organisms can respond morphologically to selection (Gould and Lewontin, 1979). For example, architectural constraints of the conserved anuran bauplan restrict the directions and magnitude of trait evolution to avoid hindering core functional performance (Lires et al., 2016; Wake, 1997). Likewise, while convergent evolution is often attributed to similar selective environments, it may also arise from shared developmental pathways and intrinsic constraints (Losos, 2011; Maynard Smith et al., 1985). However, the potential co-occurrence of bounded and OU-like dynamics inferred across traits and ecological regimes highlights a central challenge: two distinct evolutionary processes can generate statistically similar trait distributions that are, at best, difficult to distinguish using current phylogenetic comparative methods. This lack of identifiability reduces our ability to confidently infer the mechanisms underlying macroevolutionary trends. The problem is amplified in smaller clades, where limited sample sizes further reduce statistical power and manifest as ambiguous or misleading model support. Thus, these results suggest that macroevolutionary analyses can conflate adaptive and constraint-driven processes by collapsing distinct evolutionary dynamics into statistically similar signals, particularly when working with small clades that already impede model identifiability.

We hypothesized that the absence of multi-regime bounded or mixed-mode evolutionary models may bias inference toward adaptive interpretations, even in cases where the underlying trait dynamics are primarily constraint-driven. At present, model comparison frameworks in a phylogenetic context are largely restricted to contrasting multi-peak adaptive OU models (OUM) and fully unconstrained Brownian motion models with rate heterogeneity (BMS), which together represent only a limited subset of plausible evolutionary scenarios. To evaluate this shortcoming, we extracted individual sub-trees and refit single-regime models that included bounded BM and OU1, subsequently comparing their relative support to the best-fitting clade-wide OUMV model. One potential limitation of our approach is that ecologically defined regimes are not necessarily mono-phyletic, and so extracting regime-based sub-trees results in the inclusion of branch lengths over which the generating process may not have occurred (c.f. O’Meara et al., 2006). However, our validation tests of this pruning approach produced striking results that further demonstrate how adaptive and constraint-based evolutionary processes are frequently confounded due to fundamental limitations in standard comparative methods. Although the signal of dynamic adaptation was reliably detected as support for an OU1 model over unbounded or bounded BM in more species-rich regimes, tests of the species-poor ecological regimes (aquatic, semi-aquatic, torrential, burrowing, Fig. 2) frequently returned high support for bounded evolutionary dynamics over adaptive or unbounded dynamics across all three traits. Additionally, we recovered the counterintu-itive result of a ‘Goldilocks zone’ of OU1 identifiability: increasing phylogenetic half-lives initially increased support for a true OU1 model, despite this representing weaker selection, yet this pattern reverses when the half life exceeds one tenth of total tree depth (Table 2). This result contrasts previous simulation studies in which increasing *α* increases OU1 support consistently (Fig. 5C. (Boucher and Démery, 2016)). We hypothesize that when the phylogenetic half-life is too short relative to clade age, the directional component of trait evolution is rapidly erased, leaving only stochastic fluctuations around the optimum and reducing the distinctiveness of the OU1 signal. To contrast, long half-lives correspond to weakly constrained, diffusive OU1 processes in which the dynamics resemble those of Brownian motion. At a minimum, these findings warrant caution in interpreting model support as evidence for specific evolutionary mechanisms, particularly in smaller clades or putative evolutionary regimes. While it is well appreciated that comparative methods are data hungry (Burin et al., 2023) and that small sample sizes fundamentally limit the extent of conclusions that can be drawn from their results (Beaulieu et al., 2012), we suggest that these results are also reflective of a many-to-one mapping of evolutionary process to observed pattern in comparative data, analogous to the widely acknowledged many-to-one mapping of organismal form to function (Arnold, 1983). In this framework, interactions between adaptive and constraining evolutionary processes can—and do—generate statistically indistinguishable macroevolutionary patterns. Fundamentally, these results underscore the need for more flexible multi-regime modeling frameworks capable of accommodating bounded and mixed evolutionary processes, which remain underdeveloped within phylogenetic comparative methodology.

The finding that the OU1 model is consistently detectable in large but never small sub-trees is supported by our second set of simulations, which focused on single-regime bounded and OU processes across a range of appropriate tree sizes and generative parameter values. Although Boucher and Démery (2016) indicated that OU and bounded BM models can be discriminated as small tree sizes, our results differ dramatically. We found that at the smallest tree sizes, a generative OU1 model is never favored, irrespective of the strength of the rate of adaptation specified, while a true bounded model is consistently identifiable, regardless of tree size or parameter strength. Collectively, these results demonstrate a potential systematic bias toward interpreting trait evolution in small clades as being dominated by non-adaptive evolutionary constraint, even in cases where adaptive processes may be equally or more relevant. This raises a broader methodological question regarding the choice of null model in comparative analyses: when focusing on low phylogenetic signal processes, should constraint-dominated evolution be treated as the default expectation? This assumption may be justified by the ability of non-adaptive processes to generate what are typically viewed as signatures of adaptation (Gould and Lewontin, 1979; Losos, 2011). For instance, convergent evolution can arise through shared genetic architecture, correlated responses to selection acting on other traits, or even as a result of stochastic coincidence (see (Losos, 2011) for a detailed discussion). More generally, if the incorporation of constraint-focused models into comparative frameworks systematically obscures adaptive signal, it becomes important to consider which alternative evolutionary processes or model classes may be similarly overlooked under current inference approaches. Machine learning-based approaches may offer a promising avenue for addressing challenges of model identifiability by moving beyond strict likelihood-based frameworks and enabling the use of flexible, high-dimensional sets of summary statistics for classifying trait evolution under alternative generative models.

### Accepting the complex process-pattern mapping may reveal novel insights into anuran trait evolution and beyond

Accepting a many-to-one mapping between evolutionary process and observed pattern implies that adaptation operates in tandem with multiple forms of constraint in shaping functional trait evolution. Given the methodological limitations outlined above, however, it remains difficult to disentangle the relative contributions of these underlying mechanisms from trait data alone. Nevertheless, several inferences can be drawn in the context of anuran biology.

We argue that bounded dynamics provide an appropriate framework for describing body-size evolution because multiple environmental, developmental, and physiological constraints operate across clades and taxonomic scales (Brown et al., 2004; Graham et al., 1995; Hanken and Wake, 1993; Harrison et al., 2010; Levy and Heald, 2016; Peters, 1983; Rayner, 1988; Schmidt-Nielsen, 1984; Wake, 1982), potentially preventing the emergence of a single fitness optimum that is the target of selection. A range of factors are postulated to determine bound placement across biological systems (Brown et al., 2004; Emerson, 1978; Gomes et al., 2009; Gomez-Mestre et al., 2012; Levy and Heald, 2016; Newman and Dunham, 1994; White et al., 2006; Womack and Bell, 2020; Zug, 1972; Zug and Altig, 1978), however, it is difficult to evaluate their relative contributions and how these may change throughout a clade’s history. We first propose that the lower bound on body size observed in arboreal taxa is linked to constraints on the evolution of pedal morphology. These taxa exhibit specialized toe disc apparatus for surface adhesion and agility (Emerson and Diehl, 1980), including enlarged toe pads and extensive pedal webbing. Below a certain size, developing such complex structures may be precluded (Hanken and Wake, 1993), suggesting that minimum body size reflects the point at which functional digit morphology can no longer be maintained. Additionally, previous work supports systematic variation in locomotor demands within arboreal habitats, which can be subdivided into open, low, and high canopy zones (Gomes et al., 2009). If fitness optima differ across these vertical strata, aggregating all arboreal taxa may obscure underlying multi-peak dynamic adaptation, instead creating an apparent pattern of bounded trait evolution. Alternatively, the observed pattern may reflect a broad fitness plateau, within which focal trait variation has minimal impact on performance. Under this scenario, trait values may drift within bounds without strong directional selection toward a distinct optimum. More broadly, our results indicate that evolutionary mode may vary across ecological regimes for a single trait. Body size evolution is best described by an OU model in terrestrial settings but by a bounded model in arboreal settings. While previous work on anuran functional trait evolution has examined the influence of microhabitat on estimated evolutionary parameters (e.g. (Citadini et al., 2018; Gomes et al., 2009; Moen et al., 2013; Womack and Bell, 2020)) few studies have considered shifts in evolutionary mode itself. Shifts in mode imply that the balance between adaptive processes and constraints may shift at finer ecological scales than typically considered, reflecting the distinct functional challenges imposed by different habitats.

What is less well understood is how ecological shifts and fluctuating constraints across ontogeny influence the development and expression of functional traits. In lineages exhibiting the ancestral biphasic life cycle, which is characterized by free-living and feeding aquatic larvae with terrestrial adults, residual larval structures and functions may impose developmental and physiological constraints on adult morphology, leaving detectable signatures in trait data (Alberch, 1989; Hanken and Wake, 1993; Wake, 1982). However, the adaptive decoupling hypothesis (Moran, 1994) proposes that selection acting in larval and adult environments may be antagonistic, such that the constraints operating at each life stage can become partially independent (Sherratt et al., 2017). In this framework, a free-living larval stage may reduce carry-over constraints on adult morphology by allowing greater separation between larval and adult functional demands. When ecological pressures are strongly divergent between life stages, selection may favor compromise phenotypes that maintain functionality across both stages (Bonett et al., 2022), whereas more similar larval and adult environments may permit a broader range of evolutionary trajectories (Gould and Lewontin, 1979). More generally, the condensed period of developmental system restructuring that occurs during metamorphosis (Alberch, 1989) likely impacts which constraints most strongly shape morphological evolution (Bardua et al., 2021; Fabre et al., 2020; Moran, 1994). Evidence of such decoupling is observed in instances of many-to-one mappings between form and function (Arnold, 1983) across ontogeny, where diverse larval morphologies converge on similar adult phenotypes (Bragg and Bragg, 1958; Pfennig, 1990). Taken as a whole, these patterns suggest that adopting a reproductive mode-focused classification for evolutionary regimes may better capture the dynamic interplay between developmental constraints and ecological shifts across ontogeny, and provide a more mechanistic framework for understanding functional trait evolution across Anura.

The comparative methods toolkit is continuously expanding, yet methods to infer a bounded mode of evolution currently exist only in single-regime form (Boucher and Démery, 2016; Revell, 2024), which limits the degree to which signals of constraint and adaptation can be teased apart across evolutionary regimes. This limitation becomes particularly consequential in light of model non-identifiability, as patterns recovered from clade-level analyses can reflect either multi-peak adaptive evolution or primarily constraint-driven processes that produce statistically similar trait distributions. As our results suggest, multi-regime OU support may not uniquely indicate evolution on a multi-peak adaptive landscape but can also arise from bounded evolutionary dynamics operating under structural or developmental constraints (Gould and Lewontin, 1979; Lires et al., 2016; Wake, 1997). Similarly, convergent patterns attributed to shared selective regimes may instead reflect intrinsic constraints or shared developmental architecture (Losos, 2011; Maynard Smith et al., 1985). In this study, we attempted to overcome this shortfall by fitting bounded, Brownian, and OU-like models of trait evolution to individual microhabitat regimes and comparing results to clade-wide analyses. However, our simulation results further highlight the identifiability problem: even when data are generated under known multi-peak adaptive processes, regime-level inference frequently supports bounded dynamics, particularly in small clades. This mismatch underscores that adaptive and constraint-based processes can generate indistinguishable macroevolutionary signatures under current model classes, particularly when sample sizes limit statistical power.

## Conclusions

Our understanding of how generative evolutionary processes interact to produce observed patterns in trait data remains incomplete, and the remnants of Gould and Lewontin’s Panglossian paradigm remain clear. Across simulations and empirical analyses we show that constraint-based and adaptive processes can generate highly similar statistical signatures, leading to issues with model identifiability. This issue is especially pronounced in small clades, where OU-like, Brownian, and bounded processes become difficult to distinguish, and where model selection may systematically favor constraint-based interpretations even when adaptive regimes underlie trait evolution. These findings raise a wider methodological question regarding the role of null models in comparative analyses: specifically, whether constraint-dominated evolution should be treated as a default expectation in low phylogenetic signal contexts. This question is particularly relevant given that non-adaptive processes can generate patterns often interpreted as evidence of adaptation, including convergence and regime-like clustering of traits (Losos, 2011). More generally, our results suggest that current comparative frameworks may systematically ignore alternative generative mechanisms that produce indistinguishable macroevolutionary patterns. The advent of the bounded Brownian motion model represents a vital step forward in expanding the comparative toolkit, improving our ability to explicitly represent constrained evolutionary dynamics and partially addressing gaps in model identifiability. Looking ahead, approaches that extend beyond strict likelihood-based model comparison may further help alleviate these limitations. Flexible, machine learning methods that integrate high-dimensional summary statistics offer one promising avenue for discriminating among competing evolutionary processes when traditional comparative models converge on similar predictions.

## Acknowledgments

We thank David Jablonski, David Blackburn and members of the Slater Lab at The University of Chicago for thoughtful feedback on earlier versions of this manuscript. We thank David Blackburn specifically for his knowledge of anuran taxonomy. We thank Luke Harmon for constructive feedback on a later version of this manuscript and advice regarding the pruning procedure. Finally, we thank Bailey Howell at The University of Chicago Research Computing Center (RCC) for guidance on running analyses.

## Statement of Authorship

M.J. (Conceptualization, Formal analysis, Investigation, Methodology, Software, Data Curation, Writing—original draft, Writing—review & editing)

G.J.S. (Conceptualization, Writing—review & editing)

Both authors reviewed and edited the writing at all stages of revision.

