## Supplemental material for "Beyond the Panglossian paradigm: adaptation, constraint, and anuran functional trait evolution"

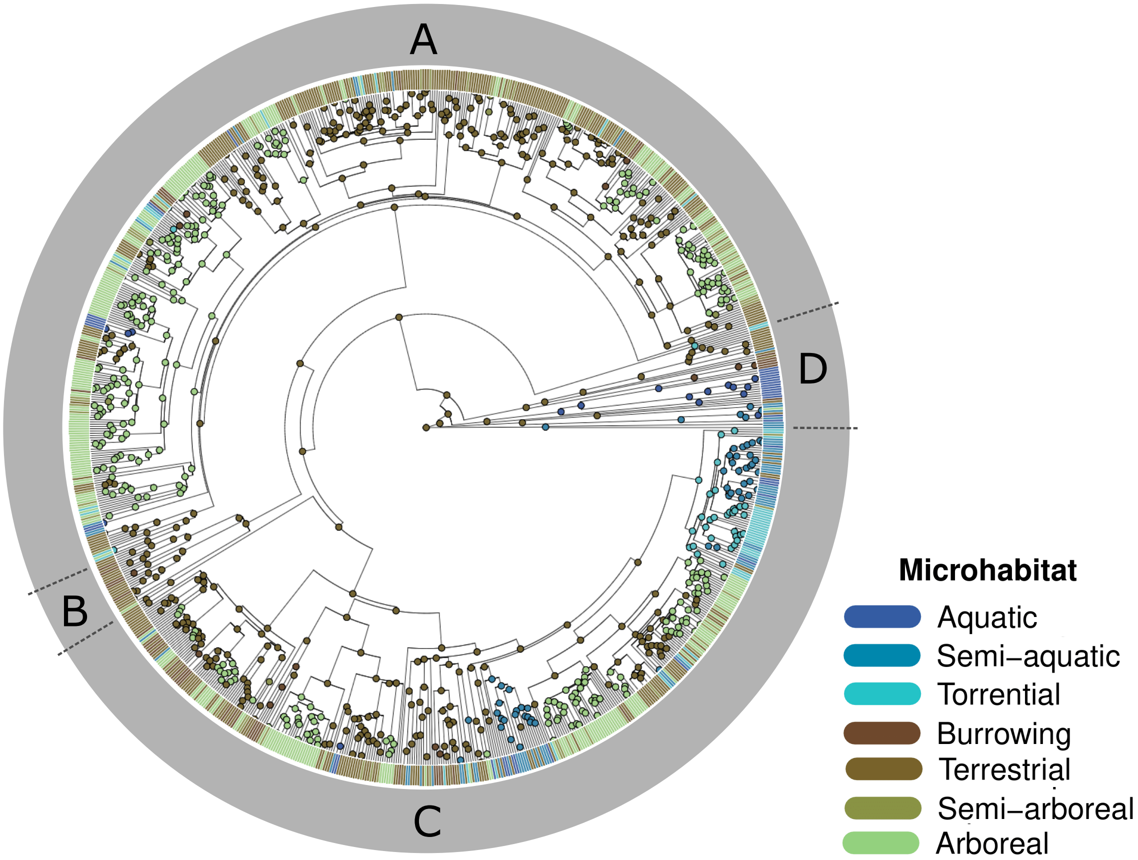


Figure S1: Maximum likelihood mapping of microhabitat states across a 927-species anuran phylogeny. States are estimated under the symmetric rates model (SYM) in ace(). Major clades are indicated as A) Hyloidea, B) Myobatrachoidea, Sooglossidae, C) Ranoidea, D) Discoglossoidea, Leiopelmatoidea, Pelobatoidea, Heleophrynidae and Pipoidea.

| | Aquatic | Semi-aquatic | Torrential | Burrowing | Terrestrial | Semi-arboreal | Arboreal |
|---|---|---|---|---|---|---|---|
| Aquatic | 0.00 | 4.10 | 3.21 | 3.09 | 7.95 | 4.00 | 7.38 |
| Semi-aquatic | 7.47 | 0.00 | 4.54 | 3.68 | 13.94 | 5.87 | 8.32 |
| Torrential | 3.50 | 6.53 | 0.00 | 3.96 | 9.85 | 5.51 | 9.03 |
| Burrowing | 2.57 | 3.67 | 3.35 | 0.00 | 9.77 | 3.86 | 7.85 |
| Terrestrial | 18.9 | 29.5 | 22.3 | 23.8 | 0.00 | 34.1 | 72.3 |
| Semi-arboreal | 3.38 | 4.89 | 4.00 | 3.75 | 13.1 | 0.00 | 13.0 |
| Arboreal | 15.2 | 19.7 | 16.6 | 19.9 | 64.4 | 32.2 | 0.00 |

Table S1: Average transition counts between habitat states. One hundred stochastic mappings were generated under the ER model using the `phytools` :: `make.simmap`() function (Revell,2024) and transition counts were averaged across them.

Table S2: Empirical support for BM, bounded, and OU1 candidate models within each pruned microhabitat-functional trait pairing. Left column labels indicate the trait; rows give support values ($W$) per microhabitat, with the greatest support values indicated in bold.

| | **Microhabitat** | **BM** | **Bounded** | **OU1** |
|---|---|---|---|---|
| **Hindlimb length** | Aquatic | 0.0276 | **0.815** | 0.157 |
| | Semi-aquatic | **0.497** | 0.294 | 0.208 |
| | Torrential | 0.302 | 0.220 | **0.478** |
| | Burrowing | 0.0115 | **0.927** | 0.0612 |
| | Terrestrial | $2.32 \times 10^{-8}$ | $1.14 \times 10^{-6}$ | **1.000** |
| | Semi-arboreal | 0.263 | **0.640** | 0.0970 |
| | Arboreal | 0.00555 | 0.00784 | **0.987** |
| **Toe webbing** | Aquatic | $3.50 \times 10^{-7}$ | **1.000** | $4.08 \times 10^{-6}$ |
| | Semi-aquatic | $4.70 \times 10^{-9}$ | **1.000** | $3.18 \times 10^{-9}$ |
| | Torrential | $5.60 \times 10^{-6}$ | **1.000** | $5.18 \times 10^{-6}$ |
| | Burrowing | 0.0217 | **0.936** | 0.0422 |
| | Terrestrial | 0.00318 | **0.955** | 0.0418 |
| | Semi-arboreal | 0.0306 | **0.958** | 0.0116 |
| | Arboreal | 0.0519 | **0.929** | 0.0191 |
| **Body size** | Aquatic | 0.0251 | 0.0408 | **0.934** |
| | Semi-aquatic | $1.93 \times 10^{-5}$ | 0.0521 | **0.948** |
| | Torrential | 0.0212 | **0.599** | 0.380 |
| | Burrowing | 0.0224 | **0.702** | 0.275 |
| | Terrestrial | $2.26 \times 10^{-7}$ | $1.03 \times 10^{-5}$ | **1.000** |
| | Semi-arboreal | 0.105 | 0.134 | **0.761** |
| | Arboreal | 0.0200 | **0.915** | 0.0652 |
